# The quail as an avian model system: its genome provides insights into social behaviour, seasonal biology and infectious disease response

**DOI:** 10.1101/575332

**Authors:** Katrina M Morris, Matthew M Hindle, Simon Boitard, David W Burt, Angela F Danner, Lel Eory, Heather L Forrest, David Gourichon, Jerome Gros, LaDeana Hillier, Thierry Jaffredo, Hanane Khoury, Rusty Lansford, Christine Leterrier, Andrew Loudon, Andrew S Mason, Simone L Meddle, Francis Minvielle, Patrick Minx, Frédérique Pitel, J Patrick Seiler, Tsuyoshi Shimmura, Chad Tomlinson, Alain Vignal, Robert G Webster, Takashi Yoshimura, Wesley C Warren, Jacqueline Smith

## Abstract

The Japanese quail (*Coturnix japonica*) is a popular domestic poultry species and an increasingly significant model species in avian developmental, behavioural and disease research. We have produced a high-quality quail genome sequence, spanning 0.93 Gb assigned to 33 chromosomes. In terms of contiguity, assembly statistics, gene content and chromosomal organization, the quail genome shows high similarity to the chicken genome. We demonstrate the utility of this genome through three diverse applications. First, we identify selection signatures and candidate genes associated with social behaviour in the quail genome, an important agricultural and domestication trait. Second, we investigate the effects and interaction of photoperiod and temperature on the transcriptome of the quail medial basal hypothalamus, revealing key mechanisms of photoperiodism. Finally, we investigate the response of quail to H5N1 influenza infection. In quail lung, many critical immune genes and pathways were downregulated, and this may be key to the susceptibility of quail to H5N1. This genome will facilitate further research into diverse research questions using the quail as a model avian species.

## INTRODUCTION

Japanese quail (*Coturnix japonica*) is a migratory bird indigenous to East Asia and is a popular domestic poultry species raised for meat and eggs in Asia and Europe. Quail have been used in genetics research since 1940 (Shimakura 1940), and are an increasingly important model in developmental biology, behaviour and biomedical studies (Minvielle 2009). Quail belong to the same family as chickens (Phasianidae) but have several advantages over chickens as a research model. They are small and easy to raise, have a rapid growth rate and a short life cycle, becoming sexually mature only seven to eight weeks after hatching (Huss et al. 2008).

Quail have become a key model in several research fields (Cheng et al. 2010). The avian embryo has long been a popular model for studying developmental biology due to the accessibility of the embryo, which permits fate mapping studies (Le Douarin and Barq 1969; Le Douarin and Kalcheim 1999) and dynamic imaging of embryogenesis (Huss et al. 2015; Bénazéraf et al. 2017; Sato et al. 2017). Several transgenic lines that express fluorescent proteins now exist, which greatly facilitates time-lapse imaging and tissue transplantation (Scott and Lois 2005; Sato et al 2010; Huss et al 2015; Moreau et al. 2018; Huss et al In Press)

The quail embryo survives manipulation and culture better than chicken embryos making them ideal for this type of research (Huss et al. 2008). Quail have been used as a model for stem cell differentiation, for example a culture system that mimics the development of hematopoietic stem cells has been recently developed, as quail show greater cell multiplication in these cultures than chickens (Yvernogeau et al. 2016). Quail are also used to study the genetics underlying social behaviours (Mills et al. 1997), sexual behaviour (Adkins-Regan 2009; Meddle et al. 1997), pre- and post-natal stress programming (Marasco et al. 2016), and emotional reactivity (Mills and Faure 1991; Jones and Mills 1999; Beaumont et al. 2005; Recoquillay et al. 2013). Japanese quail have a fast and reliable reproductive response to increased photoperiod, making them an important model species for investigation into seasonal behaviour and reproduction in birds (Robinson and Follett 1982; Nakane and Yoshimura 2010; Nakane and Yoshimura 2014). The molecular mechanisms behind seasonality including metabolism and growth, immunity, reproduction, behaviour and feather moult is poorly understood despite its importance in the management of avian species.

Quail are also important in disease research (Baer et al 2015). Different strains of quail have been developed as models of human disease such as albinism (Homma et al. 1968) or necrotizing enterocolitis in neonates (Waligora-Dupriet et al. 2009). Quail lines have also been selected on their immunological response (Watanabe and Nagayama 1979). There are key differences in the immunogenetics of quail and chicken - particularly in the major histocompatibility complex (MHC) (Shiina et al 2004; Hosomichi et al. 2006). Investigating the immunology of quail is important for understanding infectious disease spread and control in poultry. For example they are an important species for influenza transmission, with previous research showing that quail may play a key role as an intermediate host in evolution of avian influenza (Makarova et al. 2003; Perez et al. 2003; Wan and Perez 2006). Zoonotic H5N1 influenza strains have crossed from quail to human causing mortality in the past (Guan et al. 2002; Webster et al. 2002), making them a potential pandemic source.

We have produced a high-quality annotated genome of the Japanese quail (*Coturnix japonica*), and herein describe the assembly and annotation of the quail genome and demonstrate key uses of the genome in immunogenetics, disease, seasonality and behavioural research demonstrating its utility as an avian model species.

## RESULTS

### Genome assembly and annotation

We sequenced a male *Coturnix japonica* individual from an inbred quail line using an Illumina HiSeq 2500 instrument. Total sequence genome input coverage of Illumina reads was ~73x, using a genome size estimate of 1.1 Gb. Additionally, 20x coverage of long PacBio reads were sequenced and used to close gaps. The assembled male genome *Coturnix japonica* 2.0 is made up of a total of 2,531 scaffolds (including single contigs with no scaffold association) with an N50 scaffold length of 2.9 Mb (N50 contig length is 511 kb). The assembly sequence size is 0.927 Gb with only 1.7 % (16 Mb) not assigned to 33 total chromosomes. *Coturnix japonica* 2.0 assembly metrics were comparable to previous assemblies of Galliformes, and superior to other quail genomes (Oldeschulte et al. 2017; Wu et al. 2018) in ungapped (contigs) sequence length metrics (**Table 1**). Specifically, in comparison to recently published genomic data from the Japanese quail (Wu et al. 2018), our genome is substantially less fragmented (contig N50 of 0.511 Mb vs 0.027 Mb), has been assigned to more chromosomes, and has more complete annotation with ncRNA, mRNA and pseudogenes predicted. Our estimate of total interspersed repetitive elements was 19% genome-wide based on masking with Windowmasker (Morgulis et al. 2018). In the genomes of other quail species the estimated repeat content was much lower; ~10% less in both species (Oldeschulte et al. 2017).

**Table 1.** Representative assembly metrics for sequenced Galliform genomes^1^.

| Common name | Assembled version | N50 contig<br>(Mb) | N50 scaffold<br>(Mb) | Total assembly<br>size (Gb) | Assembled<br>chromosomes |
| --- | --- | --- | --- | --- | --- |
| Japanese quail | Coturnix japonica 2.0 | 0.511 | 3.0 | 0.93 | 33 |
| Japanese quail | Wu et al. PMID:<br>29762663 | 0.027 | 1.8 | 1.01 | 34 |
| Chicken | Gallus gallus 5.0 | 2.895 | 6.3 | 1.20 | 30 |
| Scaled quail | ASM221830v1 | 0.154 | 1.0 | 1.01 | NA |
| Northern bobwhite | ASM59946v2 | 0.056 | 2.0 | 1.13 | NA |
| Turkey | Turkey 5.0 | 0.036 | 3.8 | 1.12 | 33 |

<sup>1</sup>All species-specific assembly metrics derived from the NCBI assembly archive.

To improve the quantity and quality of data used for the annotation of the genome, we sequenced RNA extracted from seven tissues sampled from the same animal used for the genome assembly. Using the same inbred animal increases the alignment rate and accuracy.

The amount of data produced for annotation from the 7 tissues is (in Gb): 18.9 in brain, 35.6 in heart, 19.3 in intestine, 27.8 in kidney, 39.0 in liver, 18.8 in lung and 34.0 in muscle. High sequencing depth was aimed for in these tissues, to help detect low expression genes including those that are tissue-specific. In total we predicted 16,057 protein-coding genes and 39,075 transcripts in the *Coturnix japonica* genome (**Table 2**). In comparison to other assembled and annotated Galliformes, transcript and protein alignments of known chicken RefSeq proteins to *Coturnix japonica* suggest the gene representation is sufficient for all analyses described herein (**Table 3**). However, we find ~1000 fewer protein coding genes in the Japanese quail than the northern bobwhite (*Colinus virginianus*) and scaled quail (*Callipepla squamata*) genomes (Oldeschulte et al. 2017). We attribute this to the use of different gene prediction algorithms, and the slightly lower assembled size of Japanese quail, 927 Mb compared to 1 Gb in other quail genomes (Oldeschulte et al. 2017; **Table 1**).

**Table 2.**
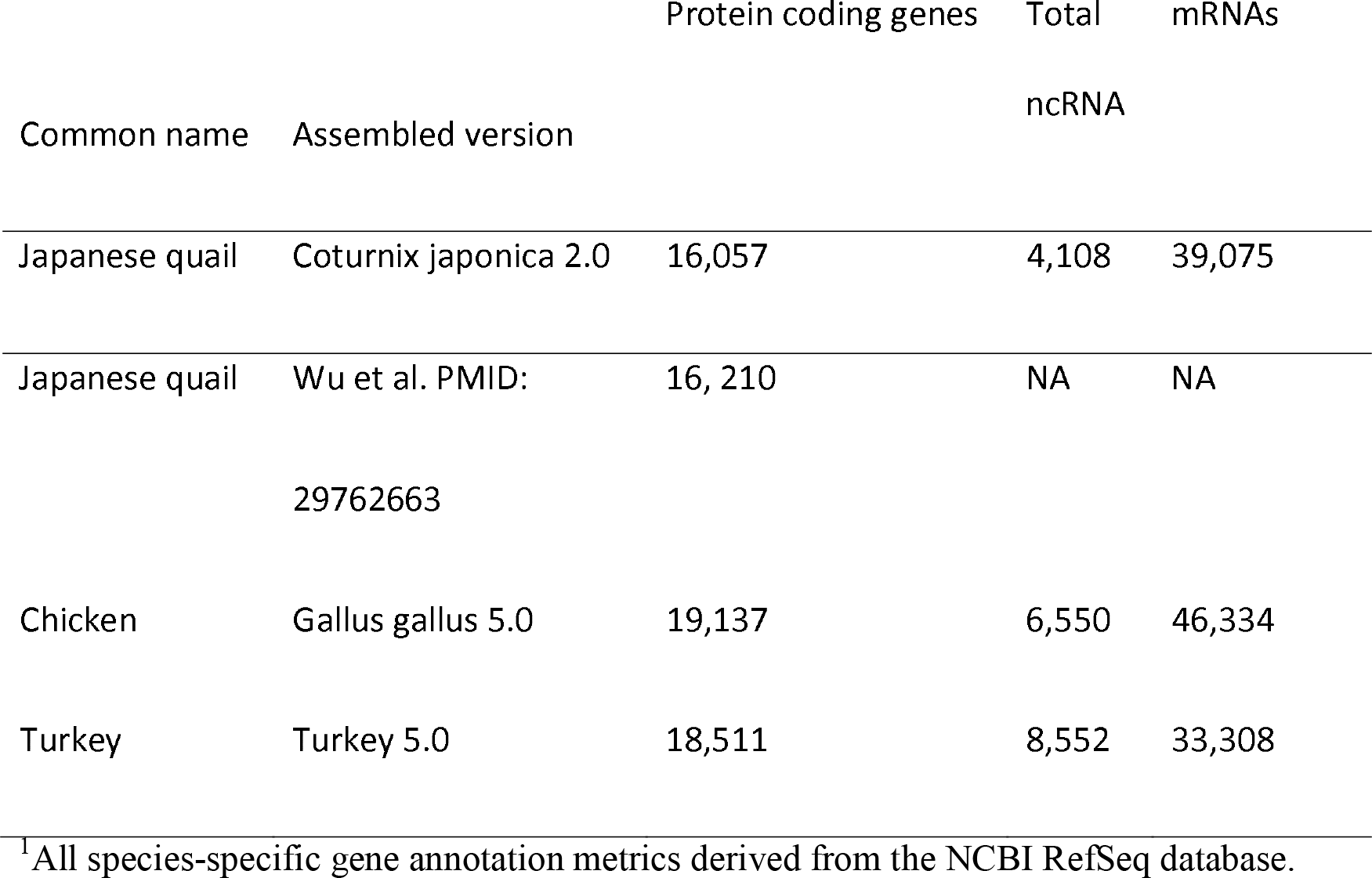
Representative gene annotation measures for assembled Galliform genomes^1^.

**Table 3.**
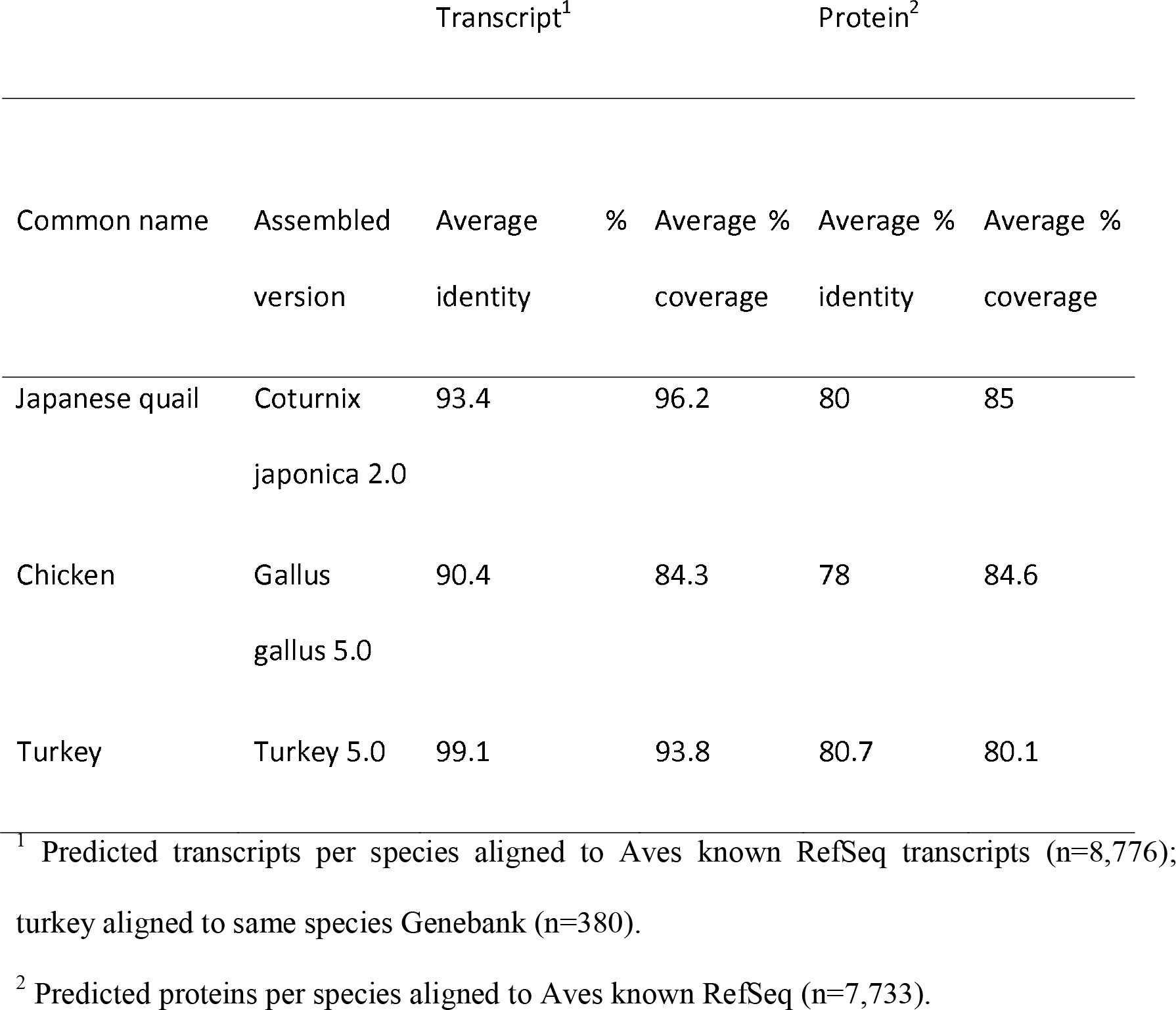
Estimates of gene and protein representation for sequenced Galliform genomes.

For further annotation, a set of genes unnamed by the automated pipeline were manually annotated. As part of an ongoing project to investigate hemogenic endothelium commitment and HSC production (Yvernogeau et al. 2016), transcriptomes were produced for two cultured cell fractions. Study of these cells is critical for developmental biology and regenerative medicine, and quail are an excellent model for studying these as they produce much more hematopoietic cells than similar chicken cultures. Approximately 8,000 genes were expressed in these cells lines which lacked gene names or annotation from the automated annotation pipeline. Using BLAST (Altschul et al. 1990) searches to identify homology to other genes, 3,119 of these were manually annotated (**Supplemental File 1**).

Genome completeness was also quantitatively assessed by analyzing 4,915 single copy, orthologous genes derived from OrthoDB v7 and v9 (Zdobnov et al. 2017). Presence and contiguity of these conserved, avian-specific genes were tested with BUSCO v3.0.2 (Waterhouse et al. 2017). A comparison with the chicken assembly (*Gallus gallus* 5.0; Warren et al. 2017) indicates that 95% of these genes are present and full length in all three assemblies. The percentage of duplicated, fragmented and missing genes are also very similar between the assemblies (**Supplemental Figure S1**). The quail genome has 10 more missing and 23 more fragmented genes than the *Gallus gallus* 5.0 assembly. However, relative to the total number of genes in the benchmarking set, these increases amount to just 0.2% and 0.5%, respectively. This indicates that the quail genome, like the chicken genome, is highly contiguous and in terms of its expected gene content, is close to complete.

### Galliforme genome synteny

Comparative mapping of the quail and chicken genomes revealed a high conservation of the chromosomal arrangement (**Fig. 1**; **Supplemental File S2**), with no major rearrangements since the divergence of the two species approximately 23 MYA (van Tuinen and Dyke 2004). All identified quail chromosomes showed synteny conservation to their chicken chromosomal counterparts. By comparison, the turkey (*Meleagris gallopavo*) genome is more highly rearranged with two chromosomes having synteny conservation to each of chicken and quail chromosomes 2 and 4 (Dalloul et al. 2010). No large intra-chromosomal translocations were seen between chicken and quail chromosomes, compared to the two seen in the turkey (Dalloul et al. 2010). Inversions and inter-chromosomal translocations were common, with 33 large (>1Mb) inversions or translocations occurring between chicken and quail chromosomes **(Fig. 2**; **Supplemental File S2**). The quail chromosomes are more compact than their chicken and turkey counterparts (14% smaller on average). This may be linked to the metabolic cost of migratory flight in quails, as previous studies have demonstrated smaller genomes and higher deletion rates in flying birds compared to flightless birds (Kapsuta et al. 2017)

**Figure 1:**
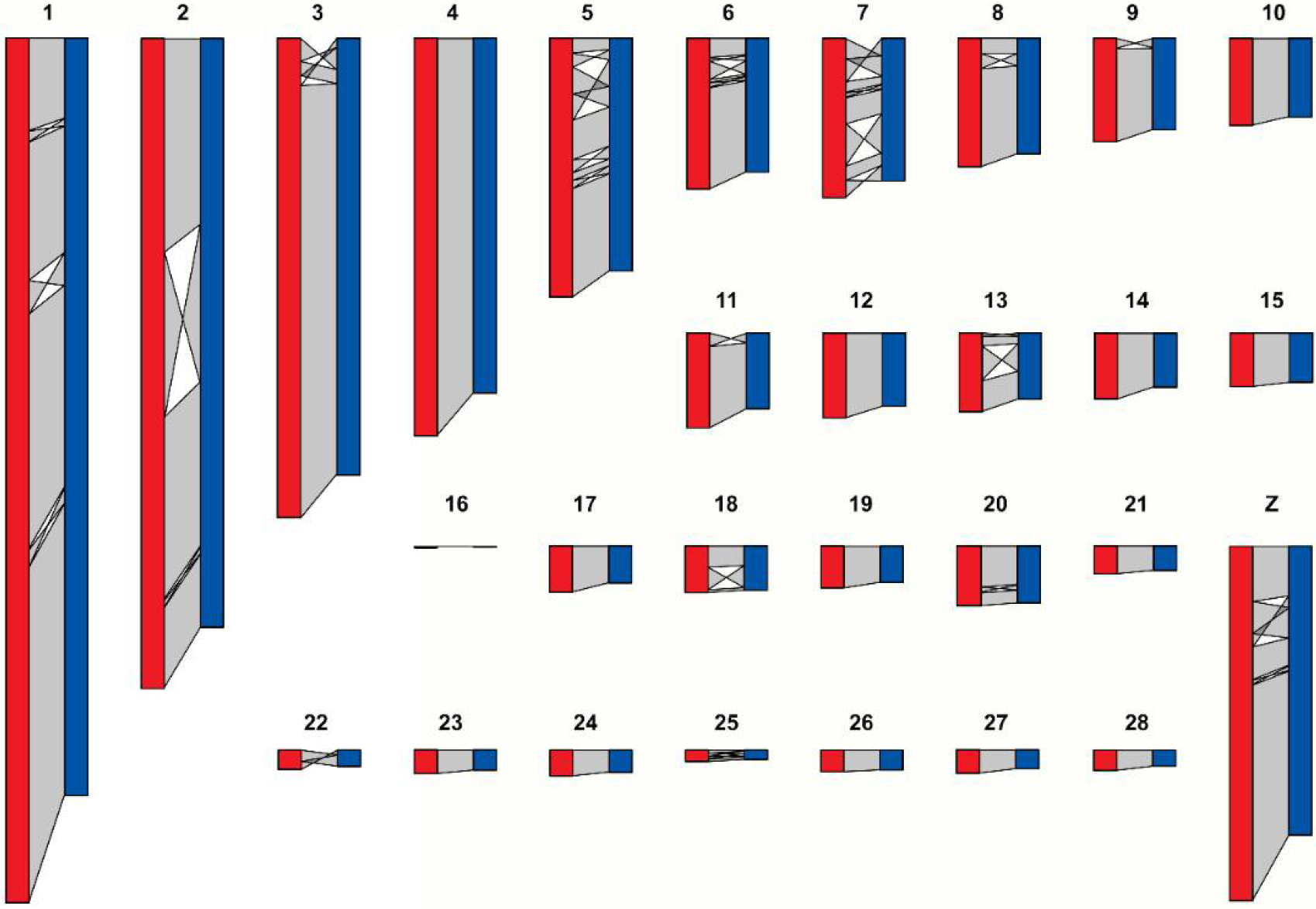
Synteny map of chicken (red) and quail (blue) chromosomes

**Figure 2:**
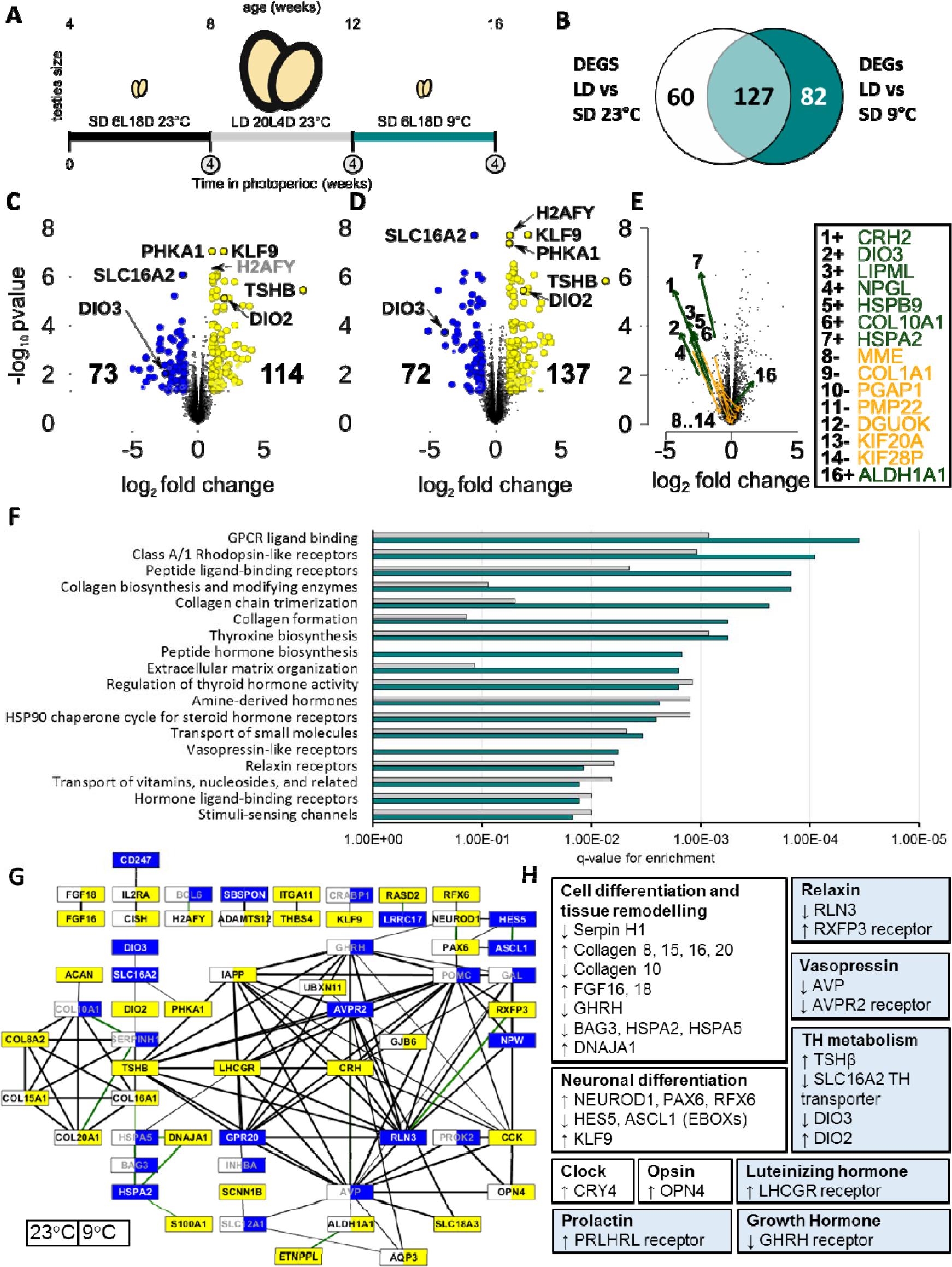
Genome-wide analysis of temperature-dependent transcriptome responses to photoperiod in quail. Experimental design showing the 3 time-points each sampled after 4 weeks of the target photoperiod (circled) with RNASeq at n=4 **A**. Intersection of DEGs between LD 23°C vs SD 23°C and LD 23°C vs SD 9°C **B**. Volcano plots comparing LD 23°C vs SD 23°C showing 71 up (yellow) and 42 down (blue) DEGs **C** and LD 23°C vs SD 23°C **D**. Grey labels do not pass fold change threshold at 23°C. Temperature-dependent effects on fold change in DEGs when comparing SD at 23°C and SD 9°C. Arrows point from 23°C to 9°C and indicate a significant amplifying (green) or dampening (orange) effect of 9°C on photoperiod response **E** significantly enriched pathways in DEG genes at LD vs SD 23°C (grey) and LD vs SD 9°C (teal) q-value thresholds **F**. Network of up (yellow), down (blue), and no significant change (white) regulated inter-connected genes (LD vs SD) using the String database. The left side of a node indicates the expression change at 23°C and right at 9°C. Edges are weighted by the combined score, and green edges represent experimental support **G**.

Orthologous genes between quail and closely related species were identified through reciprocal BLAST searches. One-to-one orthologs in chicken were identified for 78.2% of all quail genes and 91.8% of protein-coding quail genes (**Supplemental Table S1**), indicating a high degree of genic conservation in the quail genome. Fewer orthologs were seen between turkey and quail genes (69.3%), although the number of orthologs of protein-coding genes was similar (91.7%), so the discrepancy is likely due to missing non-coding gene predictions in the turkey genome. As expected, conservation of one-to-one orthologs was lower with the mallard duck (*Anas platyrhynchos*), with duck orthologs identified for 64.5% of quail genes (78.9% protein-coding genes).

### Endogenous retroviruses (ERVs)

ERVs represent retroviral integrations into the germline over millions of years and are the only Long Terminal Repeat (LTR) retrotransposons which remain in avian genomes (Mason et al. 2016; Kapusta and Suh 2017). Whilst the majority of ERVs have been degraded or epigenetically silenced, more recent integrations retain the ability to produce retroviral proteins, impacting the host immune response to novel exogneous infections (Verla et al. 2009; Aswad et al. 2012). A total of 19.4 Mb of the *Coturnix japonica* 2.0 assembly was identified as ERV sequence using the LocaTR pipeline (Mason et al. 2016; **Supplemental File S3**, **Supplemental File S4**). ERVs therefore account for 2.1% of the quail genome sequence, levels similar to those in the chicken and turkey (Warren et al. 2017; **Supplemental Table S2**), and similarly analysed passerine birds (Mason et al. 2016).

The majority of ERV sequences in all three genomes were short and fragmented, but 393 intact ERVs were identified in the quail, most of which were identified as alpha-, beta- or gamma-retroviral sequences by reverse transcriptase homology. It is possible that the smaller genome size of the quail compared to other birds reflects a more limited expansion of ERVs and other repeats (such as the LINE CR1 element; **Supplemental Table S2**) within the genome, following the basal avian lineage genome contraction (Kapusta and Suh 2017; Kapusta et al. 2017). However, ERV content is highly species-specific (Mason et al. 2016).

Despite variation in total and intact ERV content, the overall genomic ERV distribution in these three gallinaceous birds was highly similar. ERV sequence density was strongly correlated with chromosome length on the macrochromosomes and Z chromosome (r > 0.97; P < 0.001), but there was no significant correlation across the other smaller chromosomes. Furthermore, ERV density on each Z chromosome was at least 50% greater than would be expected on an autosome of equal length. These results support the depletion of repetitive elements in gene dense areas of the genome, and the persistence of insertions in poorly recombining regions, as was seen in the chicken (Mason et al. 2016). This is further supported by the presence of clusters of intact ERVs (where density was five times the genome-wide level) on the macrochromosomes and sex chromosomes (**Supplemental Table S2**).

### Immune gene repertoire

We investigated the immune genes in the quail genome in detail due to the importance of quail in disease research. The MHC-B complex of the quail has been previously sequenced and found to be generally conserved compared to chicken in terms of gene content and order (Shiina et al. 2004; Hosomichi et al. 2006). However the quail MHC contains a higher copy number of several gene families within the MHC-B (Shiina et al. 2004) and shows increased structural flexibility (Hosomichi et al. 2006), as well as an inversion in the *TAP* region (Shiina et al. 2004). The MHC-B sequence in the quail genome extends from the previously sequenced scaffold, and this additional region also contains similar gene content and order to chicken, but with gene copy number variations. As in the chicken, the *CD1A* and *B* genes are found downstream of the MHC I region, while many *TRIM* family genes and *IL4I1* are encoded upstream. The BG region, which encodes a family of butrophylin genes known as *BG* genes in the chicken, was also present in the quail. Within this region, six *BG* genes were identified in the quail, compared to thirteen in the chicken (Salomonsen et al. 2014). At least five of these *BG* genes are transcribed in the quail lung and ileum. The chicken and turkey have an additional MHC locus known as the Rfp-Y or MHC-Y locus, which contains several copies of non-classical MHCI-Y and MHCIIB-Y genes. However, no MHC-Y genes have been previously identified in quail. BLAST searches of both the quail genome and quail transcriptomes, as well as the bobwhite and scaled quail genomes, failed to identify any MHC-Y genes, indicating this locus probably does not exist in the quail.

Cathelicidins and defensins are two families of antimicrobial peptides that have activities against a broad range of pathogens and exhibit immune-modulatory effects. Orthologs of all four chicken cathelicidins and of thirteen chicken defensins (Cheng et al. 2015) were identified in the quail genome (**Supplemental File S5**). Due to their high divergence, of the thirteen defensins only four were annotated through the annotation pipeline, with the remainder identified through BLAST and HMMER searches with chicken defensins. The only poultry defensin missing from the quail genome is *AvBD7*. The defensins are encoded in a 42 kb cluster on quail chromosome 3, as in chickens. A 4 kb gap in the scaffold in this region may explain the missing *AvBD7* sequence.

Several genes are thought to be crucial for influenza resistance in both humans and birds, including *RIG-I*, *TLR* and *IFITM* genes. *RIG-I* has not previously been identified in chicken, despite being present in ducks and many other bird orders, and is considered highly likely to be deleted from the chicken genome (Barber et al. 2010). In addition, an important RIG-I binding protein RNF135 has also not been identified in chicken (Magor et al. 2013). Likewise, an ortholog of *RIG-I* or *RNF135* could not be identified in the quail genome or transcriptomes through BLAST searches and therefore is likely missing in the quail also. Orthologs of all five chicken *IFITM* genes (*IFTIM1, 2, 3, 5* and *10*) were identified in the quail genome and transcriptomes. In addition, orthologs of each chicken TLR, including key TLRs for viral recognition, *TLR4* and *TLR7*, were identified in the quail genome, except that of *TLR1A*.

### Selection for social motivation

Quail has been used as a model to study the genetic determinism of behaviour traits such as social behaviours and emotional reactivity (Beaumont et al. 2005; Recoquillay et al. 2013; Recoquillay et al. 2015), these being major factors in animal adaptation. Moreover quail selected with a low social motivation behave in a way that can be related to autistic-like traits, so the genes and causal variants are of wider interest to the biomedical community. Here we use the new quail genome assembly to improve previous results on the detection of selection signatures in lines selected for sociability. Due to the non-availability of a useable quail reference genome at the start of these studies, genomic sequence data produced from two DNA pools of 10 individuals each from two quail lines diverging for social motivation had been aligned to the chicken reference genome, GallusWU2.58 (Fariello et al. 2017). As a result, only 55% of the reads had mapped in proper pairs, whereas by using our quail genome as a reference, this number increased to 92%. This corresponds to an improvement of the averaged coverage from 9x to 20x and of the number of analysed SNPs from 12,364,867 to 13,506,139.

The FLK (Bonhomme et al. 2010) and local (Fariello et al. 2017) score analysis led to the detection of 32 significant selection signature regions (*p* < 0.05), (**Supplemental File S6**). **Supplemental Figure S2** shows an example of such a region on Chr20. This represents a substantial improvement in the number of detected regions, compared with the 10 regions obtained when using the chicken genome as a reference (Fariello et al. 2017). Of the 32 detected regions, six may be merged in pairs due to their physical proximity, four regions map to new linkage groups absent in the previous analysis, and eight correspond with results obtained in the previous study (**Supplemental File S6**). Altogether, 17 new regions were detected. Of these, eight could be seen in the previous analysis, but had not been considered as they did not reach the significance threshold, and nine are solely due to the availability of our quail assembly. Two very short selection signatures previously detected using the chicken assembly as reference are not recovered here and were most probably false positives.

These results confirm the selection signature regions harbouring genes involved in human autistic disorders or being related to social behaviour (Fariello et al. 2017; *PTPRE*, *ARL13B*, *IMPK*, *CTNNA2*). Among the genes localised in the newly detected genomic regions, several have also been shown to be implicated in autism spectrum disorders or synaptogenic activity (**Supplemental File S6**): mutations in the *EEF1A2* gene (Eukaryotic elongation factor 1, alpha-2) have been discovered in patients with autistic behaviours (Nakajima et al. 2015); *EHMT1* (Euchromatin Histone Methyltransferase 1) is involved in autistic syndrome and social behaviour disorders in human and mouse (Nakajima et al. 2015; Kleefstra et al. 2006; Balemans et al. 2010; Mitra et al. 2017); *LRRTM4* (Leucine Rich Repeat Transmembrane Neuronal 4) is a synapse organizing protein, member of the *LRRTM* family, involved in mechanisms underlying experience-dependent synaptic plasticity (Roppongi et al. 2017).

Autistic spectrum disorders are observed in several disorders that have very different aetiology, including fragile X Syndrome, Rett Syndrome or Foetal Anticonvulsant Syndrome. While these disorders have very different underlying etiologies, they share common qualitative behavioural abnormalities in domains particularly relevant for social behaviours such as language, communication and social interaction (Rutter 1978; American Psychiatric Association 2000). In line with this, several experiments conducted on high social (HSR) and low social (LSR) reinstatement behaviour quail indicate that the selection program carried out with these lines is not limited to selection on a single response, social reinstatement, but affect more generally the ability of the quail to process social information (Marasco et al. 2016). Differences in social motivation, but also individual recognition have been described between LSR and HSR quail (Balemans et al. 2010; Mitra et al. 2017). Inter-individual distances are longer in LSR quail (Balemans et al 2010) and LSR young quail have decreased interest in unfamiliar birds (Roppongi et al. 2017) and lower isolation distress than HSR ones (Recoquillay et al. 2013).

Further experiments will be required to examine the possible functional link between the selected genes and the divergent phenotype observed in these lines. Also, by analyses of genes known to be differentially expressed in the zebra finch during song learning we hope to comparatively understand molecular systems linked to behaviour in the avian brain.

### A model for avian seasonal biology

Quail is an important model for studying seasonal biology. Seminal work in quail established that pineal melatonin (Ralph et al. 1967; Lynch 1971) is regulated by the circadian clock (Cockrem and Follett 1985). In mammals, photo-sensing is dependent on a single retinal photoreceptor melanopsin (*OPN4*) that regulates pineal melatonin release. Nocturnal melatonin is critical for mammalian neuroendocrine response to photoperiod and is likely to target melatonin receptors in the *pars tubularis* (PT; Wood and Loudon 2014). Birds have a distinct non-retinal mechanism for photoreception through deep-brain photoreceptors (Menaker 1968) and melatonin does not appear to be critical for most avian seasonal cycles (Yoshimura 2013). The medial basal hypothalamus (MBH) seems to be a critical region for avian perception of photoperiod (Yasuo et al. 2003). There are currently three main candidates for avian deep-brain photoreceptors that communicate the photoperiod signal to seasonal cycles: *OPN4* (Haas et al. 2017), neuropsin (*OPN5*; Nakane et al. 2010) and vertebrate ancient (*VA;* García-Fernández et al. 2015).

While melatonin may not be a critical component to avian photoperiod signal transduction it may play a role. Photoperiodic regulation of Gonadotropin-inhibitory hormone (GnIH), first identified in quail, has been shown to be regulated by melatonin (Chowdhury et al. 2010). Melatonin receptors are also located in the quail PT (Cozzi et al. 1993) and like the mammalian PT (Lincoln et al. 2002) the expression of core clock genes in the quail PT (Yasuo et al. 2004) are phase-shifted with photoperiod. Previously, two studies (Yasuo et al. 2003; Ikegami et al. 2015) have examined temperature dependent effects of photoperiod on core clock genes, *TSHβ* in the PT and *DIO2* and *DIO3* in the MBH. Here, we leverage the new quail genome for genome-wide analysis to determine how photoperiod and temperature interact to determine the MBH transcriptome (**Fig. 2A**).

We examined the effect of short-(SD) and long-day (LD) photoperiod (SD, 6L18D & LD, 20L4D) and temperature (9°C and 23°C) (**Fig. 2A; Supplemental Figure S3**) on genome-wide transcription and identified 269 significantly differentially expressed genes (DEGs; FDR<0.05, log^2^FC>1; **Supplemental File S7**). 127 DEGs were regulated irrespective of temperature, 60 and 82 DEGs were specific to the contrast with SD 9°C and 23°C, respectively.

We identified 16 temperature dependent DEGs with a large modulating effect of temperature (log^2^FC>1) (**Fig. 2E**). With the exception of aldehyde dehydrogenase (*ALDH1A1*), the temperature-dependent photoperiod effected DEGs were down-regulated in LD. There was an equal division of genes between temperature dependent amplification and suppression of LD down-regulated genes.

The MBH shows strong TSHβ induction in LD (**Fig. 2C-D**, log^2^FC=7.96 at 9°C, 8.36 at 23°C), indicating the stamp contains the adjacent PT as well as the MBH. Ikegami et al. (Ikegama et al. 2015) *in-situ* data support the localisation of TSHβ in the quail PT. Consistent with previous MBH findings (Ikegami et al. 2015), we observed significant up-regulation of *DIO2* and down-regulation of *DIO3*, in LD. We also observed a significant effect of cold (9°C) in short days as an amplifier of *DIO3* LP down-regulation (**Fig. 2E,** log^2^FC=-3.86 at 9°C, −2.51 at 23°C). We were unable to confirm any significant effect of cold on *DIO2*. We note significant photoperiod-dependent down-regulation of the thyroid hormone specific transporter *SLC16A2* in LP that was amplified at 9°C (log^2^FC=-1.19 at 9°C, −1.63 at 23°C).

Differential regulation of G-protein coupled receptor (GPCR) signalling was the most enriched pathway regulated by photoperiod (**Fig. 2F**; **Supplemental File S8**). It also emerged as the largest connecting component within the String interaction network of DEG genes (**Fig. 2G**). TSHβ itself binds to the GPCR THR (Millar et al. 2012). G-protein signalling is also critical for opsin signalling (Shichida et al. 2009). We also observed transcriptional regulation in other GPCR hormone receptors, including Relaxin, Vasopressin, LH, Prolactin, and GH. GnRH is associated with VA opsins in AVT neurones and has been suggested as a photoperiod sensor (García-Fernández et al. 2015). We also noted down-regulation of the neuronally important GPCR GPR20 (**Fig. 2G**). In mice, deficiency of GPR20 is associated with hyperactivity and may play a role in cAMP-dependent mitogenesis (Hase et al. 2008). There was a strong enrichment of collagen biosynthetic processes and extracellular matrix organisation processes (**Fig. 2F)** and a large body of genes associated with cell differentiation and development (**Fig. 2H**).

We observed photoperiod-dependent regulation of a single clock gene, *CRY4*. *CRY4* is up-regulated in LP (log^2^FC=0.85 at 23°C, 1.37 at 9°C). This is consistent with the finding of Yasuo et al. (Yasuo et al. 2003), that the expression of *PER2-3*, *CLOCK*, *BMAL1*, *CRY1-2* and *E4BP4* remain stable across photoperiods. Intriguingly, CRY4 has recently been proposed as a component of light-dependent magneto-reception in the avian retina (Pinzon-Rodriguez et al. 2018).

We detected photoperiod effects on *OPN4* transcripts, which were up-regulated in LD. Photoperiod-dependent expression in *OPN4* may well play a role in the photoperiod-refractory response. Encephalopsin (*OPN3*) was found to be highly expressed in the MBH (2.31..2.42 log^2^CPM) but without significant changes in expression. OPN3 has recently been identified in the hypothalamus of chick hatchlings (Kato et al. 2016) but not as yet to the MBH of adult birds. *OPN5* (−0.46..-0.89 log^2^CPM) and *VA* (−0.11..0.31 log^2^CPM) were also unchanging and expressed at a low level in the MBH sample.

In conclusion, we confirm the importance of temperature and photoperiod-dependent regulation of thyroid hormone metabolism in the avian MBH (**Fig. 3**). Temperature-dependent amplification and suppression of the photoperiod response may indicate qualitative differences in the MBH pathways or simply reflect different stages of progression through seasonally phased processes. This could be further investigated by contrasting across time series at different temperatures. We also observed concurrent regulation of multiple hormonal signalling pathways, this may reflect a diversity of pathways and cell types in the MBH or reflect a corrective mechanism to account for cross-talk with other GPCR pathways. We observed LH, PRL and GH receptor transcript changes which may indicate modulation of a GnRH-anterior pituitary feedback mechanism. Intriguingly, we also note LD induction of CRY4 which raises the question - is there a role for seasonally active magneto-sensing in the MBH? In addition to observing high *OPN3* expression in the MBH, we also noted LD overexpression of *OPN4*, which could provide a potential component for an avian photoperiod-refractory mechanism. This study has demonstrated the utility of genome-wide transcriptome analysis in quail to provide valuable insights and novel hypotheses for avian seasonal biology.

**Figure 3:**
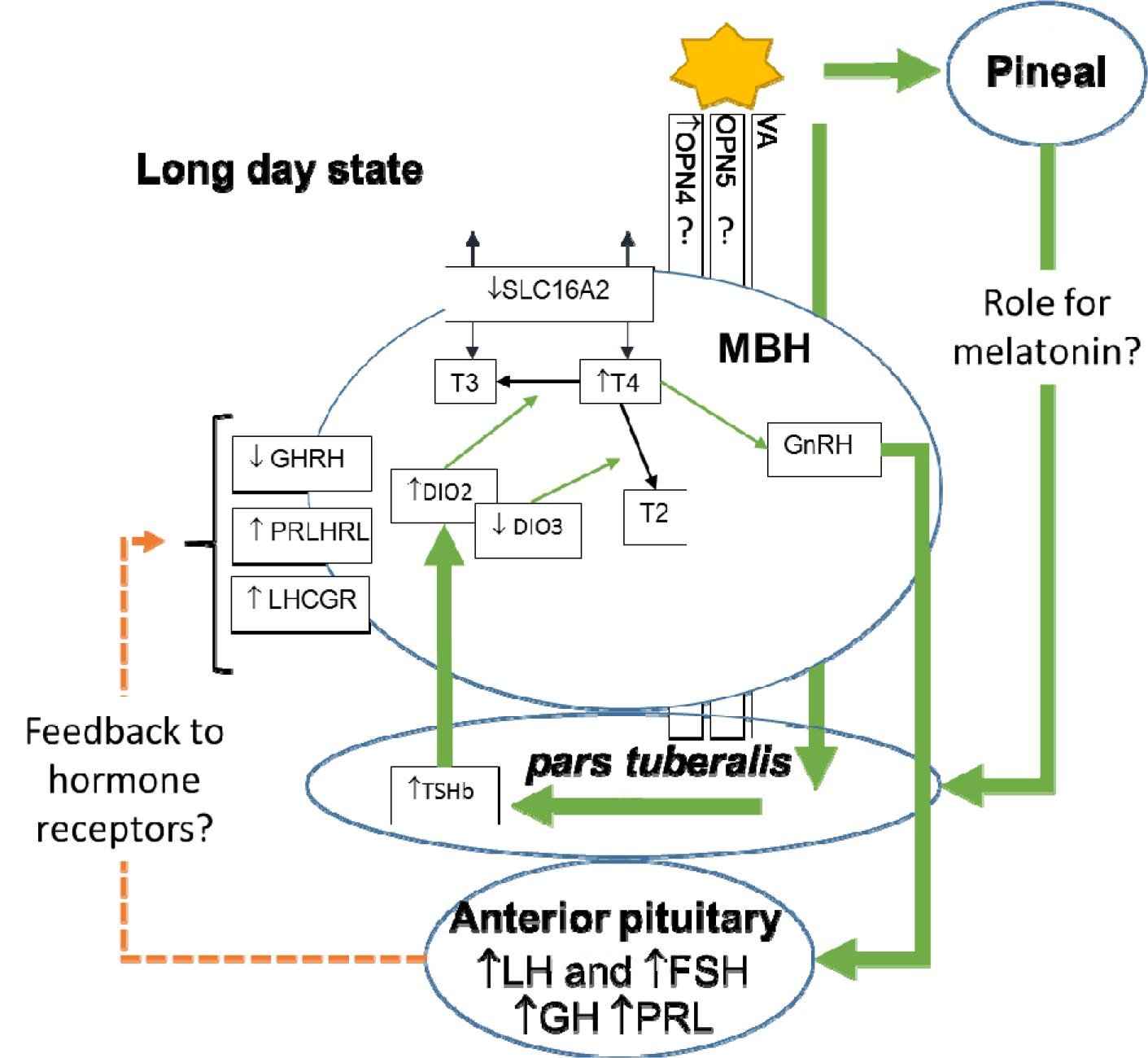
Photoperiod signalling in the MBH incorporating observations from RNASeq

### Quail response to H5N1 influenza

Highly pathogenic influenza A viruses (HPAI), such as H5N1, are responsible for enormous economic losses in the poultry industry and pose a serious threat to public health. While quail can survive infection with low pathogenic influenza viruses (LPAI), they experience high mortality when infected with strains of HPAI (Bertran et al. 2013). Quail are more susceptible than chickens to infection by some strains of H5N1 including one that caused human mortality (Webster et al. 2002). Previous research has shown that quail may play a key role as an intermediate host in the evolution of avian influenza, allowing viral strains to spread from wild birds to chickens and mammals (Makarova et al. 2003; Perez et al. 2003; Webster et al. 2002; Nguyen et al. 2016). Unlike quail and chicken, aquatic reservoir species such as duck are tolerant of most HPAI strains (Cornelissen et al. 2013). The generation of a high-quality quail genome has enabled us to perform a differential transcriptomic analysis of gene expression in quail infected with LPAI and HPAI, to better understand the response of quail to influenza infection. Lung and ileum samples were collected at 1 day post infection (1dpi) and 3 days post infection (3dpi). We also reanalysed previous data collected from duck and chickens (Smith et al. 2015) and compare this to the quail response.

To provide an overview of the response to LPAI and HPAI in quail we examined pathway and GO term enrichment of DEGs (see **Supplemental File S9, Supplemental File S10 and Supplemental Figures S4-7**). In response to LPAI infection, pathways enriched in the ileum included metabolism, JAK/STAT signalling, IL6 signalling and regulation of T-cells (**Supplemental Figure S4)**. In the lung, pathways upregulated included complement, IL8 signalling, and leukocyte activation (**Supplemental Figure S5)**. In the lung at 3dpi highly enriched GO terms included ‘response to interferon-gamma’, ‘regulation of NF-kappaB‘, ‘granulocyte chemotaxis’ and ‘response to virus’ (**Supplemental Figure S6)**, which are key influenza responses. This indicates an active immune response occurs to LPAI infection in quail, involving both ileum and lung, but with the strongest immune response occurring in the lung.

Genes upregulated in response to HPAI in the ileum were related to metabolism and transport, while inflammatory response was downregulated at 1dpi (**Supplemental Figure S6**). Downregulated pathways at 1dpi included IL-6, IL-9 and neuro-inflammation signalling pathways (**Supplemental Figure S6**). In the quail lung many genes were downregulated after HPAI infection (**Supplemental File S9**). At 3dpi, most downregulated pathways and terms were linked to immune system processes. GO terms with the highest fold enrichment in downregulated genes at this time included T and B cell proliferation, TNF signalling pathway, TLR pathway and IFN-G production **(Supplemental File S10**). Pathways downregulated included both Th1 and Th2 pathways, T cell, B cell and macrophage signalling pathways **(Supplemental Figure S7**). This indicates that crucial immune responses in quail, are downregulated in ileum, and particularly in the lung at day 3, following HPAI infection.

To compare the response of quail, duck and chicken, clustering of gene counts was examined using BioLayout 3D (Theocharidis et al. 2009). This revealed a cluster of 189 genes that were strongly upregulated at 1dpi in the duck, which showed no response in chicken and quail (**Supplemental Table S3**). This cluster was dominated by RIG-I pathway and IFN response genes including *IFNG*, *DDX60*, *DHX58*, *IRF1*, *IRF2*, and *MX1*. Pathways associated with this cluster includes MHCI processing and death receptor signalling (**Fig. 4**). Thus, the lack of this early anti-viral response may be key to the susceptibility of Galliformes to HPAI.

**Figure 4:**
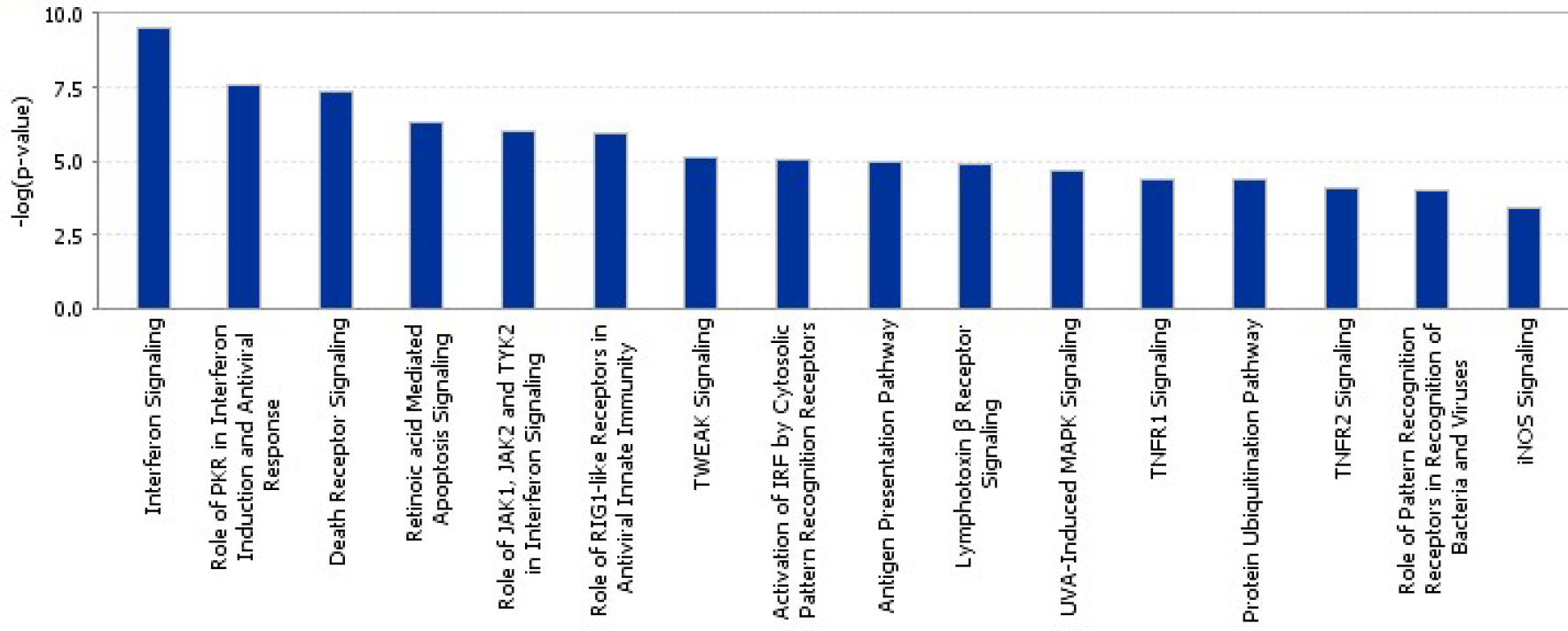
Enriched pathways in a cluster of genes highly expressed in duck lung after HPAI infection

To further compare the responses between the three species, enrichment of pathways in each species was examined (**Fig. 5**). This revealed very few commonly regulated pathways between the three species. However, at 1dpi in the ileum and 3dpi in the lung there were many pathways that were downregulated in the quail, not altered in chicken, and upregulated in the duck. In the ileum at 1dpi, this included pattern recognition and death receptor signalling. In the lung at 3dpi this involved host of immune related pathways including production of NOS by macrophages, pattern recognition, B and T cell signalling and NK-KB, IL8 and IL2 signalling.

**Figure 5:**
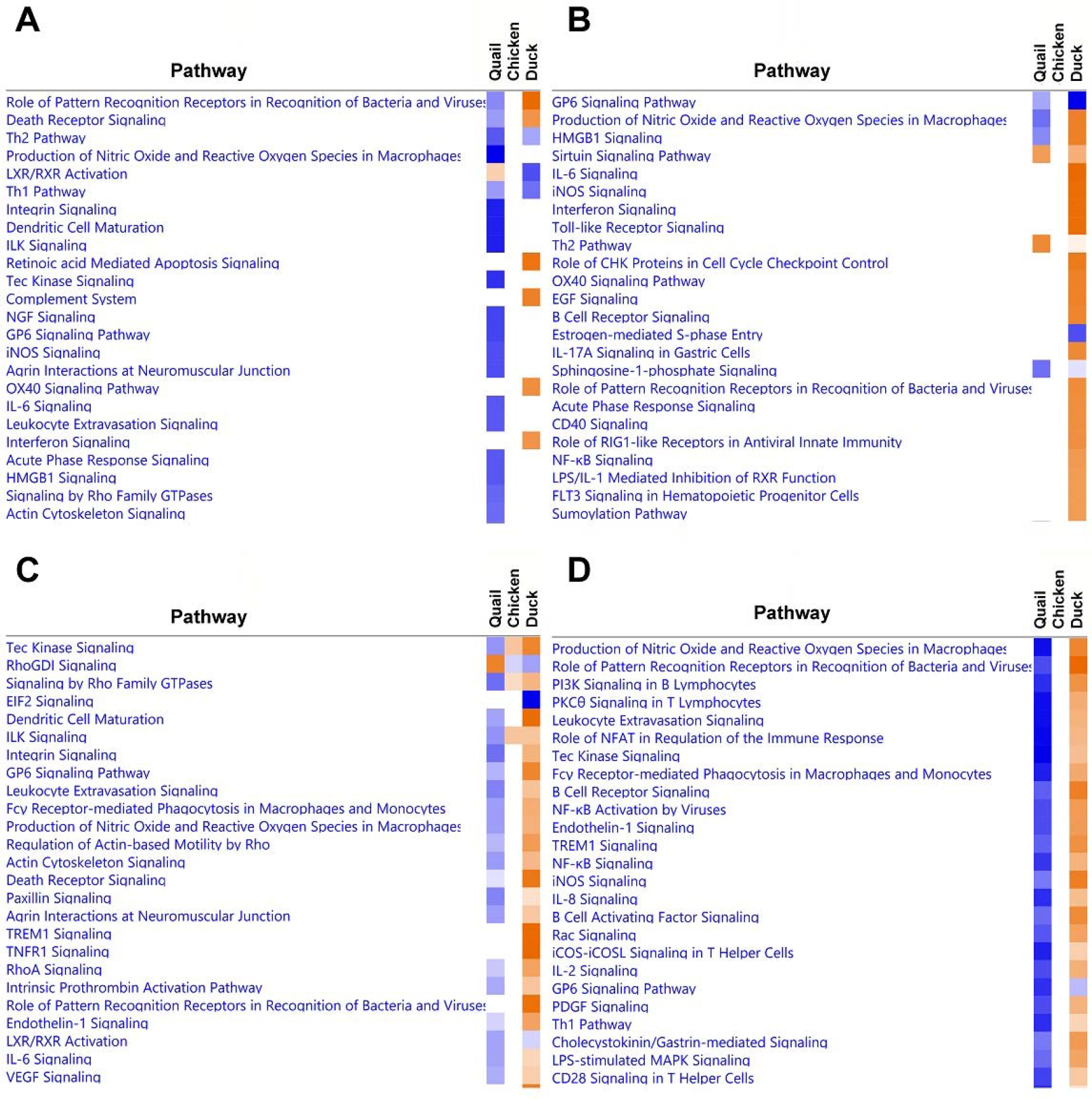
Heat-map comparison between pathways upregulated (orange) and downregulated (orange) in quail, chicken and duck, in ileum day 1 (A), ileum day 3 (B), lung day 1 (C) and lung day 3 (D).

The proportion of genes commonly regulated between quail, chicken and duck to HPAI infection was also examined (**Fig. 6**). Consistent with the heatmap comparison (**Fig. 5**), the response of chicken, quail and duck were largely unique, with few genes commonly differentially expressed. There was a large set of genes that were upregulated in duck, while being downregulated in quail at 3dpi, in both ileum and lung. In lung these genes were related primarily to innate immune system pathways, including pattern recognition pathways, cytokine production, leukocyte adhesion, TNF production, interferon production, B cell signalling and response to virus (**Supplemental File S10)**. Genes with the greatest differential expression included *RSAD2* which inhibits viruses including influenza, *IFIT5* which senses viral RNA and *OASL* which has antiviral activity. These differences further highlight that the anti-viral immune response is dysregulated in quail. Additionally in both ileum and lung the apoptosis pathway was enriched in duck, but not quail (**Supplemental File S10**). Apoptosis is known to be a critical difference in the response of chickens and ducks to HPAI infection (Kuchipudi et al. 2010).

**Figure 6:**
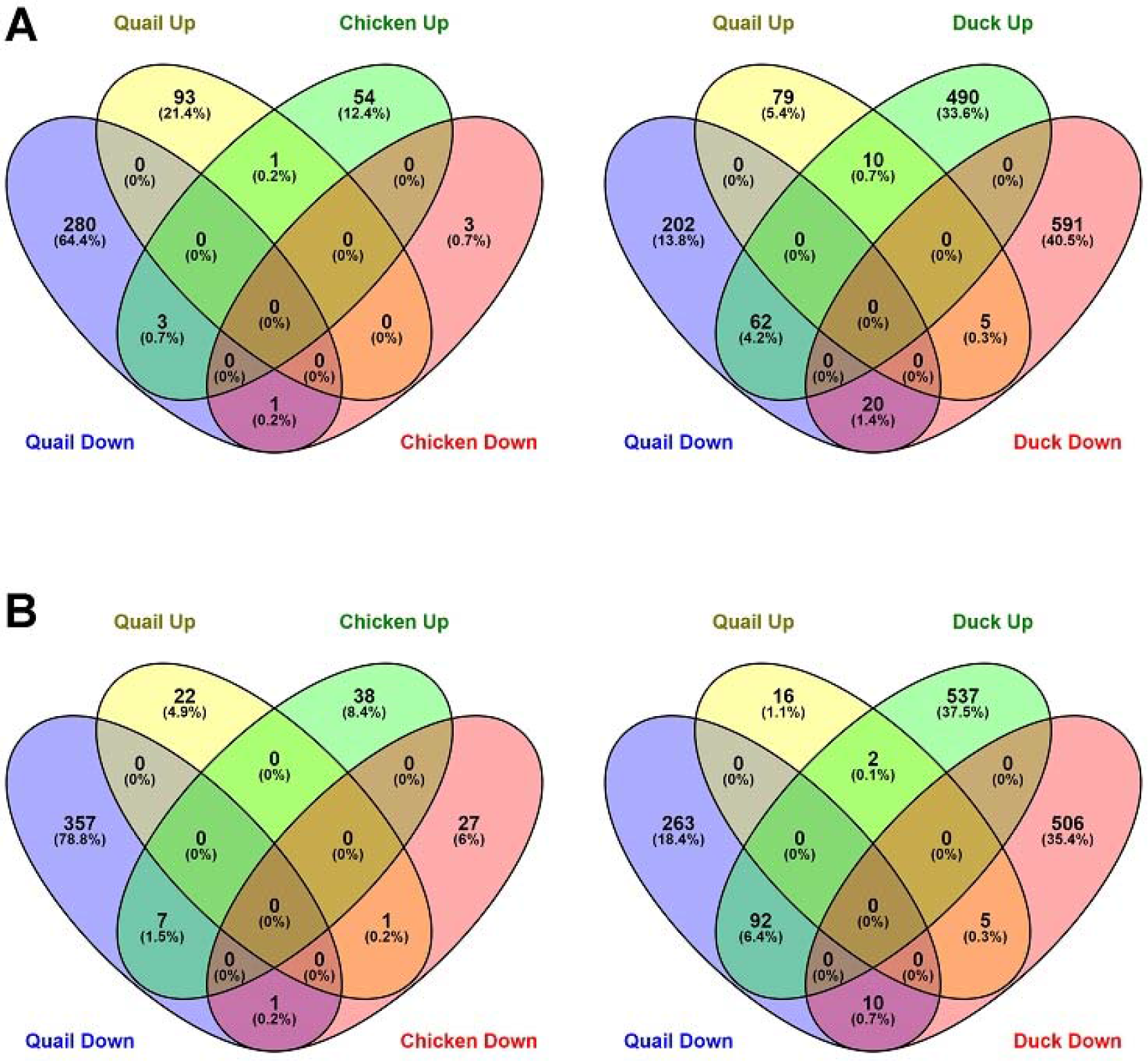
Proportion of genes commonly regulated between quail and chicken or duck to H5N1 infection on day 3 in ileum (A) and lung (B)

Lastly, we examined the response of key families involved in influenza and immune response, focussing on the lung (**Supplemental Table S4 and Supplemental File S11**). *IFTIM* genes have previously been found to have a crucial role in HPAI resistance (Smith et al. 2015) and may block AIV from entering cells (Amini-Bavil-Olyaee et al. 2013). Consistent with previous findings in the chicken (Smith et al. 2015), quail showed no significant upregulation of *IFTIM* genes, while these genes in duck were strongly upregulated, (**Supplemental Table S4**). TLRs and MHC receptors are involved in recognition of foreign molecules and triggering either an innate (TLR) or adaptive (MHC) immune response. TLR3, 4 and 7, which bind viral RNAs, were upregulated in response to LPAI in quail. A reversal was seen in response to HPAI, with *TLR4*, and *7* substantially downregulated. Likewise, genes of both MHC class I and II were upregulated in response to LPAI and downregulated in response to HPAI. By comparison there was no perturbation of TLR and MHC genes in chicken and upregulation of class I genes in duck. The quail seems to have a highly dysfunctional response to HPAI infection with key innate and adaptive immune markers downregulated at 3dpi, which contrasts with the strong immune response mounted by the duck and minimal immune response in the chicken.

In conclusion, we found that quail have a robust immune response to infection with LPAI, allowing them to survive the infection. However, they show dysregulation of the immune response after infection with HPAI, and this may explain their susceptibility to HPAI strains. IFITM response was not seen against HPAI while genes associated with apoptosis were downregulated, potentially allowing the virus to easily enter cells and spread early in infection. Antiviral and innate immune genes, including those involved in antigen recognition, immune system activation, and anti-viral responses were downregulated at 3dpi, which would prevent an effective immune response and viral clearance once infection is established. This study provides crucial data that can be used to understand the differing response of bird species to AIV, which will be critical for managing and mitigating these diseases in the future.

## DISCUSSION

Here we describe the assembly, annotation and use of a high-quality quail genome, an important avian model in biological and biomedical research. This genome will be crucial for future avian genome comparative and evolutionary studies, and provides essential genetic and genomic reference information, molecular information for making precise primers and nucleic acid probes, and accurate perturbation reagents including morpholinos, RNA inactivation tools, and CRISPR-Cas9 constructs. We have demonstrated the utility of this genome in both infectious disease and behavioural research providing further confirmation of the importance of quail as a research model, and for its role in agricultural and animal health studies. Specifically, the availability of this genome has allowed us to make significant discoveries in the unique response of quail to highly pathogenic avian influenza infection, helping elucidate the basis for extreme susceptibility seen in this species. It has also allowed us to identify and confirm genes and genomic regions associated with social behaviour, showing many similarities to genes associated with autism in humans and thus represents a possible biomedical model for autism. Furthermore, we have shown that genome-wide transcriptomics using this genome facilitated further insights and hypothesis into the mechanism of photo-periodism in avian seasonal biology. Moving forward, the availability of a high-quality quail genome will facilitate the study of diverse topics in both avian and human biology, including disease, behaviour, comparative genomics, seasonality and developmental biology.

## METHODS

### Whole Genome Sequencing and Assembly

To facilitate genome assembly by avoiding polymorphism, we produced an individual as inbred as possible. We started with a quail line previously selected for early egg production and having a high inbreeding coefficient (Minvielle et al. 1999) and four generations of brother-sister matings produced a dedicated line “ConsDD” (PEAT, INRA Tours, France). A 15 week-old male *Coturnix japonica* (id. 7356) was then selected from this line for the sequencing project. Genomic DNA was extracted from a blood sample using a high-salt extraction method (Roussot et al. 2003). Our sequencing plan followed the recommendations provided in the ALLPATHS2 assembler (Maccallum et al. 2009). This model requires 45x sequence coverage of each fragment (overlapping paired reads ~180 bp length) from 3 kb paired end (PE) reads as well as 5x coverage of 8 kb PE reads. These sequences were generated on the HiSeq2500 Illumina instrument. Long-reads used for gap filling were generated at 20x coverage on the same DNA source using a RSII instrument (Pacific Biosciences). The Illumina sequence reads were assembled using ALLPATHS2 software (Maccallum et al. 2009) using default parameter settings and where possible, and scaffold gaps were closed by mapping and local assembly of long-reads using PBJelly (English et al. 2012). The Illumina long insert paired-end reads (3 kb and 8kb PE) were used to further extend assembled scaffolds using SSPACE (Boetzer and Pirovano et al. 2014). The draft assembly scaffolds were then aligned to the genetic linkage map (Recoquillay et al. 2015) and the Galgal4.0 chicken reference (Genbank accession: GCA_000002315.2) to construct chromosome files following previously established methods (Warren et al. 2017). Finally, all contaminating contigs identified by NCBI filters (alignments to non-avian species at the highest BLAST score obtained), and all contigs < 200 bp were removed prior to final assembly submission.

### Gene Annotation

Specific RNA-seq data for the genome annotation was produced from the same animal used for the genome assembly. RNA was extracted from heart, kidney, lung, brain, liver, intestine, and muscle using Trizol and the Nucleospin^®^ RNA II kit (MACHEREY-NAGEL), following the manufacturer’s protocol.

The *Coturnix japonica* assembly was annotated using the NCBI pipeline, including masking of repeats prior to *ab initio* gene predictions, for evidence-supported gene model building. We utilized an extensive variety of RNA-Seq data to further improve gene model accuracy by alignment to nascent gene models that are necessary to delineate boundaries of untranslated regions as well as to identify genes not found through interspecific similarity evidence from other species. A full description of the NCBI gene annotation pipeline was previously described (Thibaud-Nissen et al. 2013). Around 8,000 lacked gene symbols from this pipeline, and these were further annotated manually by using BLAST searches using the corresponding sequences and extracting protein names from Uniprot.

### Comparative analyses

A set of single copy, orthologous, avian-specific genes were selected from OrthoDB v. 9 (Zdobnov et al. 2017) and their status (present, duplicated, fragment or missing) were tested with BUSCO v.3.0.2 (Waterhouse et al. 2017) in the *Gallus gallus* 5.0 and *Coturnix japonica* 2.0 genomes. *Ab initio* gene predictions were done within the BUSCO framework using tBLASTn matches followed by avian specific gene predictions with Augustus v. 3.3 (Stanke et al. 2006). Gene status was assessed by running HMMER (Eddy 1998) with the BUSCO HMM profiles of the orthologous sequences. Comparative maps and breakpoint data were generated using AutoGRAPH (Derrien et al. 2007) using chicken and quail gff annotation files, using default settings.

### Endogenous retrovirus identification

Endogenous retroviruses (ERVs) were identified in the *Coturnix japonica* 2.0 and Turkey 5.0 genome assemblies using the LocaTR identification pipeline (Mason et al. 2016) and compared to a previous analysis of ERVs in the Gallus gallus 5.0 genome assembly (Warren et al. 2017). LocaTR is an iterative pipeline which incorporates LTR_STRUC (McCarthy and Donald 2003), LTRharvest (Ellinghaus et al. 2008), MGEScan_LTR (Rho et al. 2007) and RepeatMasker (http://repeatmasker.org) search algorithms.

### Sociability selection study

The data and methods used have been described previously (Fariello et al. 2017). Briefly, two quail lines were used, divergently selected on their sociability (Mills and Faure 1991): high social (HSR) and low social (LSR) reinstatement behaviour. A total of 10 individuals from generation 50 of each quail line were sequenced after equimolar DNA pooling. Sequencing was performed (paired-ends, 100 bp) on a HiSeq 2000 sequencer (Illumina), using one lane per line (TruSeq sbs kit version 3). The reads (190,159,084 and 230,805,732 reads, respectively, for the HSR and LSR lines) were mapped to the CoJa2.2 genome assembly using BWA (Li and Durbin 2009), with the mem algorithm. Data are publicly available under SRA accession number SRP047364. Within each line, the frequency of the reference allele was estimated for all SNPs covered by at least 5 reads, using Pool-HMM (Boitard et al. 2013). This analysis provided 13,506,139 SNPs with allele frequency estimates in the two lines. FLK values (Bonhomme et al. 2010) were computed for all these SNPs, and the local score method (Fariello et al. 2017) was applied to the p-value on single-marker tests.

### Photoperiod study

MBH tissue was collected as previously (Ikegami et al., 2015). Male 4-week old quail were obtained from a local dealer in Japan and kept under SD conditions (6L18D) for 4 weeks. At 8 weeks of age, quail were transferred to LD conditions (20L4D) and kept under LD conditions for 4 weeks to develop their testes. And then, 12 week-old LD quail were transferred to short day and low temperature (SL: 6L18D 9C) conditions for another 4 weeks to fully regress their testes. All samples were collected at ZT18. (Light onset is same for LD and SD and light offset was extended in LD group). RNA-Seq was performed using a TruSeq stranded mRNA prep (Revision E 15031047) with 125bp paired-end reads on a HiSeq Illumina 2500 with four replicates in each of the three conditions.

Reads were quality (Phred>25) and adapter trimmed with trim galore (version 0.4.5). Tophat (version 2.1.0; Kim et al. 2013) with bowtie2 (version 2.2.6) was used to map reads to the quail genome (GCA_001577835.1 *Coturnix japonica* 2.0), using the NCBI annotation. We determined feature counts for gene loci using the featureCounts program (Liao et al. 2014) in the subread (version 1.5.0) package (Liao et al. 2013). Statistical analysis was performed using the limma package (Law et al. 2014; version 3.36.1) in the R programming environment (version 3.5.0). The trimmed mean of M-values normalization method (TMM) was used for normalisation with Voom for error estimation (Sup. Table 2). We retained gene loci with more than 10x coverage in three replicates in at least two conditions. A categorical least squared regression model was fitted using LD 23°C, SD 23°C, and SD 9°C conditions. Statistics for pairwise comparisons were then recalculated by refitting contrasts to the model for LD 23°C vs SD 23°C, LD 23°C vs SD 9°C and SD 23°C vs SD. The Benjamini Hochberg approach (Benjamini and Hochberg 1995) was used to estimate the false discovery rate. For reporting numbers of photoperiod significant genes, we applied thresholds of FDR <0.05, log2 CPM > 0, and absolute log2 fold change > 1. Temperature-dependent genes are reported as those with a photoperiod significant effect at either 23°C or 9°C and a significant effect when contrasting SD 9°C and SD 23°C at the same thresholds defined across photoperiods.

### Influenza response study

All experiments involving animals were approved by the Animal Care and Use Committee of St. Jude Children’s Research Hospital and performed in compliance with relevant policies of the National Institutes of Health and the Animal Welfare Act. All animal challenge experiments were performed in animal biosafety level 2 containment facilities for the LPAI challenges and in biosafety level 3 enhanced containment laboratories for the HPAI challenges. Viral challenges of quail, tissue collection, RNA extractions and sequencing were carried out as previously described for chicken (Smith et al. 2015). Briefly, fifteen mixed-sex quail were challenged with 106 EID50 intranasally, intratracheally, and intraocularly of LPAI A/Mallard/British Columbia/500/2005 (H5N2) in phosphate buffered saline (PBS). Fifteen quail were challenged with 101.5 EID50 intranasally, intratracheally, and intraocularly of HPAI A/Vietnam/1203/2004 (H5N1) in PBS. Mock infection control groups (*n*=12) were also inoculated, receiving an equivalent volume and route of administration with PBS. Animals were monitored daily for clinical signs. Lung and ileum samples were collected from all birds on 1dpi and 3 dpi. RNA extractions were performed using Trizol and QIAGEN’s RNeasy kit. For sequencing thirty-six cycle single-ended sequencing was carried out on the Genome Analyser IIx using Illumina v3 Sequencing by Synthesis kits.

All quail as well as duck and chicken RNA-seq reads from the previous study (Smith et al. 2015) were analysed as follows. Ileum and lung RNAs were analysed from PBS infected control (3 samples from each of 1dpi and 3dpi), H5N1-infected (3 samples from each of 1dpi and 3dpi, except quail ileum 1dpi which had 2 samples) and H5N2-infected (3 samples from each of 1dpi and 3dpi). 251 million reads of 36 nucleotides in length were generated in total for quail. Reads were quality checked using FastQC and trimmed for quality using Trim-galore. Mapping was performed to the quail genome (GCA_001577835.1 Coturnix japonica 2.0), chicken genome (GCA_000002315.3 Gallus_gallus-5.0) and duck (GCA_000355885.1 BGI_duck_1.0) using Tophat2 (Kim et al. 2013) using default options. For quantification and differential analysis, the following pipeline was used. First transcripts were assembled and quantified using cufflinks (Trapnell et al. 2010), guided with the NCBI annotation for the relevant genome, and the multi-read correct option was used. The transcriptomes were merged using stringtie merge (Pertea et al. 2015) and cuffdiff (Trapnell et al. 2010) was used for differential analysis using default settings. To determine orthology between quail, duck and chicken genes, reciprocal BLAST searches were performed. For analysis of GO term enrichment the PANTHER overrepresentation test (Thomas et al. 2003) was used and for pathway analysis Ingenuity Pathway Analysis software (QIAGEN) was used. For clustering analysis BioLayout 3D (Theocharidis et al. 2009) was used using default settings except 1.4 inflation for Markov clustering.

## DECLARATIONS

### ETHICS STATEMENT

ConsDD, HSR and LSR animals were bred at INRA, UE1295 Pôle d’Expérimentation Avicole de Tours, F-37380 Nouzilly, in accordance with European Union Guidelines for animal care, following the Council Directives 98/58/EC and 86/609/EEC. Animals were maintained under standard breeding conditions and subjected to minimal disturbance. Furthermore, the ethics committee approved the rearing protocol (authorization number 00915.02). The use of quail in photoperiod experiments were approved by the Animal Experiment Committee of Nagoya University. All experiments involving animals in the infection studies were approved by the Animal Care and Use Committee of St. Jude Children’s Research Hospital and performed in compliance with relevant policies of the National Institutes of Health and the Animal Welfare Act.

### DATA ACCESS

All data generated or analysed during this study are included in this published article (and its additional files), or in the following public repositories. Data has been submitted to the public databses under the following accession numbers: genome sequence data, NCBI Genome [GCA_001577835.1] (https://www.ncbi.nlm.nih.gov/assembly/GCF_001577835.1/); transcription annotation data, SRA [SRR2968870 - SRR2968911] (https://www.ncbi.nlm.nih.gov//bioproject/PRJNA296888); RNA-seq data for infection studies, Array Express, quail [E-MTAB-3311] (https://www.ebi.ac.uk/arrayexpress/experiments/E-MTAB-3311/), duck [E-MTAB-2909] (https://www.ebi.ac.uk/arrayexpress/experiments/E-MTAB-2909/), chicken [E-MTAB-2908] (https://www.ebi.ac.uk/arrayexpress/experiments/E-MTAB-2908/); Sequencing of HSR/LSR lines, SRA [SRP047364] (https://www.ncbi.nlm.nih.gov/bioproject/PRJNA261665); RNA-seq data for photoperiod study, SRA [PRJNA490454] (https://www.ncbi.nlm.nih.gov/bioproject/?term=PRJNA490454). This quail reference genome will be available on the Ensembl platform as part of release 96.

## ACKNOWLEDGEMENTS

The authors would like to thank the Edinburgh Genomics sequencing facility (Edinburgh, UK) for carrying out the transcriptomic sequencing, RNA sequencing of the reference individual was performed on the GeT-Plage platform (http://get.genotoul.fr/en/) and funded by the INRA Genetics Division (QuailAnnot program). HSR and LSR lines sequencing was supported by the French Agence Nationale de la Recherche (SNP-BB project, ANR-009-GENM-008). We are grateful to the genotoul bioinformatics platform Toulouse Midi-Pyrenees (Bioinfo Genotoul) for providing help, computing and storage for these resources and to the ITAVI SeqVol program for financing gallo-anseriformes genome sequencing. KMM was supported by a National Health and Medical Research Council Overseas Postdoctoral Fellowship. This work was funded in part by the National Institute of Allergy and Infectious Diseases, National Institutes of Health, under contract numbers HHSN266200700005C and HHSN272201400006C, by ALSAC and by BB/N015347/1 and an HFSP 2015 award (RGP0030/2015).

## AUTHOR CONTRIBUTIONS

Inbred quail line for sequencing: FM, DG; Selection for social motivation: CL, DG; Genome sequencing and assembly: WW, JG, CT, PM, LH, DB; Transcriptome and annotation: AV, FP, TJ, HK, RL; Assembly quality assessment; LE; ERV analysis: AM; selection signature analyses: FP, SB, AV; photoperiod study: TY, DB, TS, SM, AL, MMH; avian flu studies: RW, JPS, KM, AD, HF, JS, DB; preparation of manuscript: KM, JS, TY, SM, AL, MMH, DB, AV, WW, LE, AM, RL.

