## Supplemental Figures for "The quail as an avian model system: its genome provides insights into social behaviour, seasonal biology and infectious disease response"

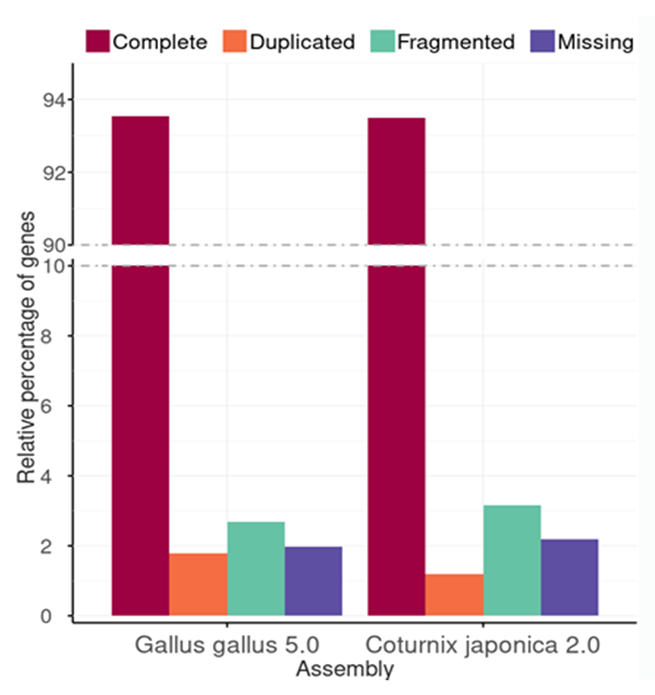

**Supplemental Figure S1: Transcriptome and assembly completeness with BUSCO 3.**

Assessment of 4,915 single copy, orthologous, avian genes in two chicken and the quail genome assemblies were done using the BUSCO v. 3.0.2 pipeline. Complete genes are single copy orthologs; duplicated appear in multiple copies; fragmented are shorter than the expectation from the BUSCO profile match; missing is not found.

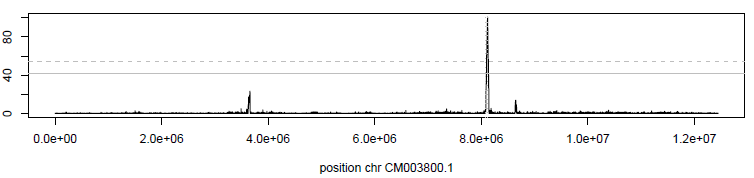

**Supplemental Figure S2: Example of selection signature on quail chromosome 20.**

The local score (Lindley process based on the score function -log10(pFLK) - 1) is shown along genomic positions on chromosome 20 (bp).

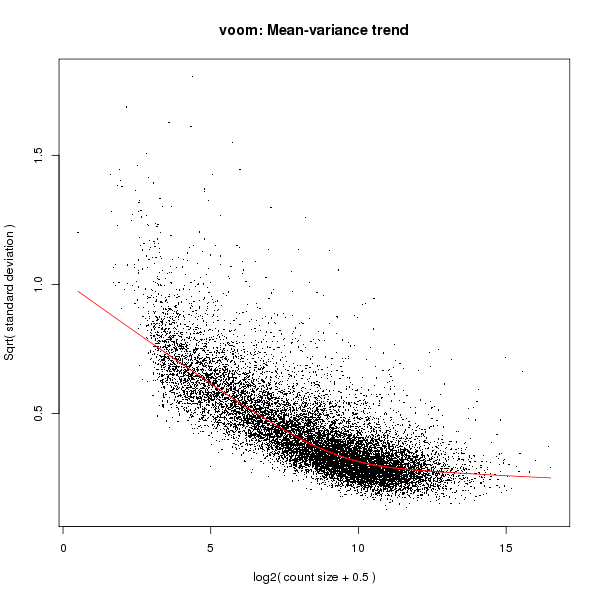

**Supplemental Figure S3**: **Voom estimation of the mean-variance trend in the RNASeq**

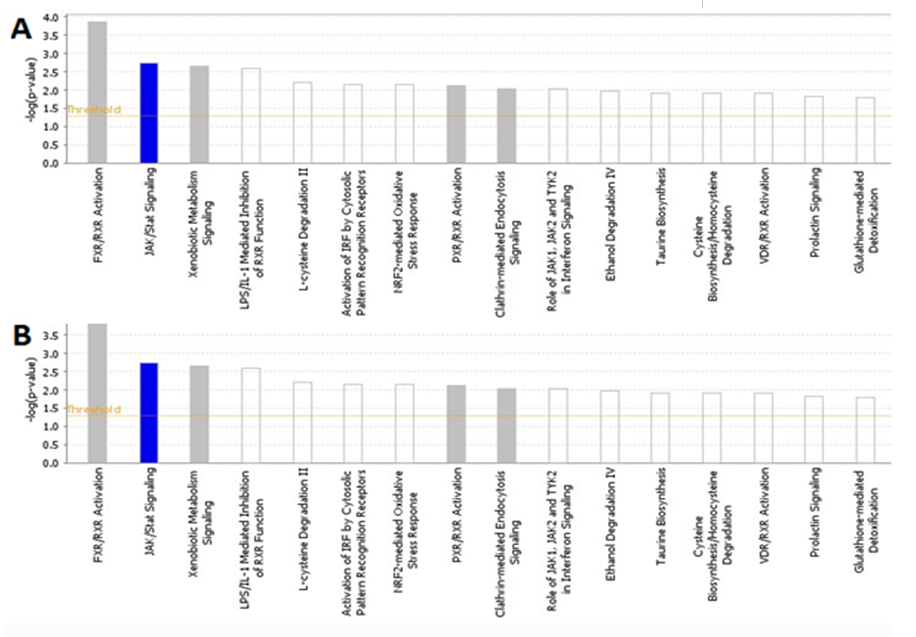

**Supplemental Figure S4:**

Biological pathways significantly altered in the ileum of LPAI infected quail vs controls (1 day post infection (A) 3 days post infection (B))

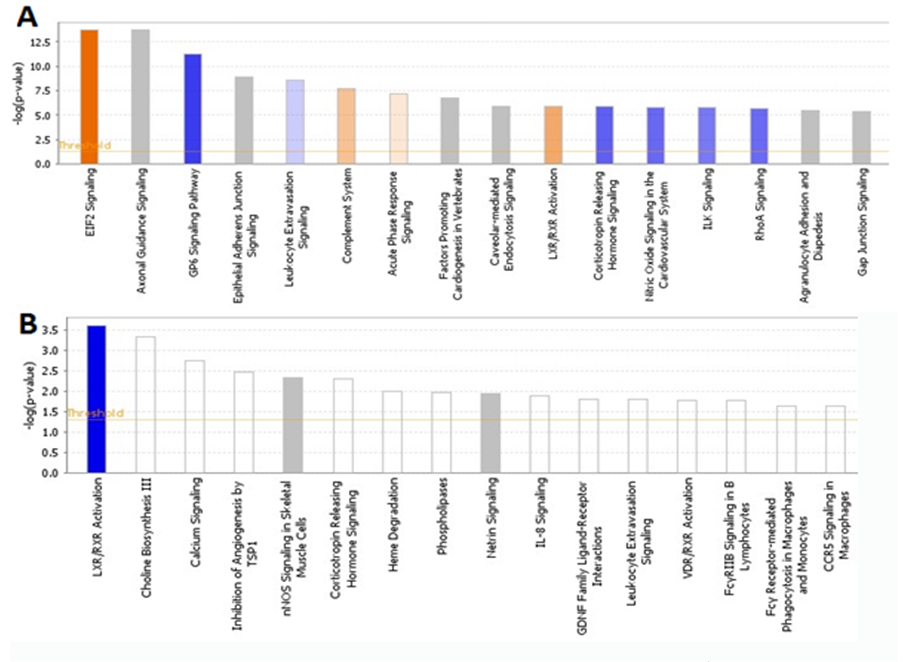

**Supplemental Figure S5:**

Biological pathways significantly altered in the lung of LPAI infected quail vs controls (1 day post infection (A) 3 days post infection (B))

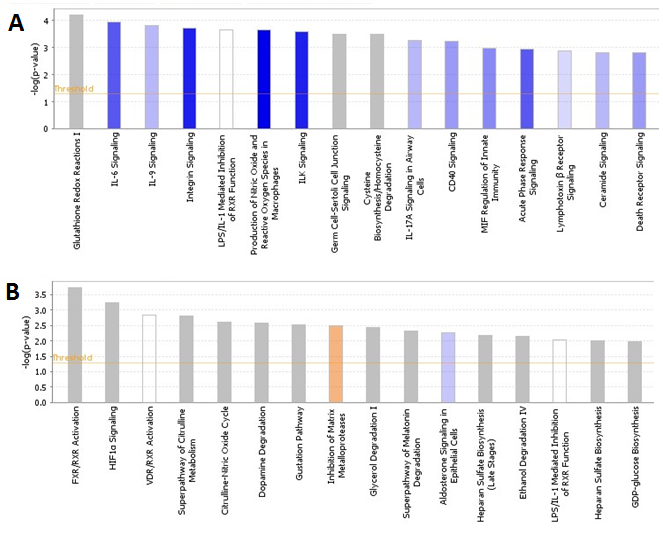

**Supplemental Figure S6:**

iological pathways significantly altered in the ileum of HPAI infected quail vs controls (1 day post infection (A) 3 days post infection (B))

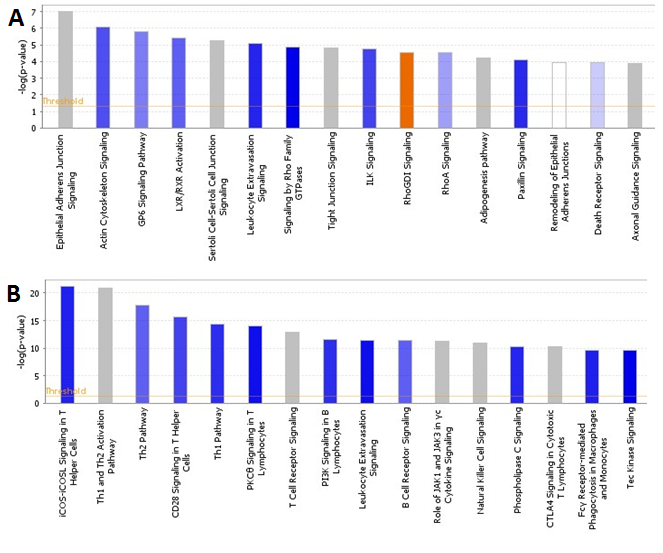
 **Supplemental Figure S7:**

Biological pathways significantly altered in the lung of HPAI infected quail vs controls (1 day post infection (A) 3 days post infection (B))

**Supplemental Table S1:** Percent of quail genes with orthologs identified in related bird genomes

|  | Chicken | Turkey | Duck |
| --- | --- | --- | --- |
| All genes | 78.2 | 69.3 | 64.5 |
| Protein-coding genes | 91.8 | 91.7 | 78.9 |

**Supplemental Table S2:** A comparative summary of assembled ERVs in quail, chicken and turkey.

|  | Coturnix japonica 2.0 | Gallus gallus 5.0 | Turkey 5.0 |
| --- | --- | --- | --- |
| Total base length (bp) | 927,656,957 | 1,230,258,557 | 1,128,339,136 |
| CR1 copy number | 184,261 | 225,230 | 252,607 |
| ERV content | 19,390,614 (2.1%) | 34,811,469 (2.8%) | 31,641,338 (2.8%) |
| Intact ERVs | 393 | 1,212 | 411 |
| Intact ERVs in clusters | 138 (35.1%) | 458 (37.8%) | 150 (36.5%) |

**Supplemental Table S3:** List of the cluster of genes upregulated in duck at day 1 during HPAI infection

| Genes | | | |
| --- | --- | --- | --- |
| ABCA8 | EPSTI1 | MITD1 | SAMD9L |
| ABCA9 | FBXO18 | MKL2 | SAMHD1 |
| ADAP2 | FBXW11 | MOV10 | SARAF |
| ADAR | FGF7 | MX1 | SCARB2 |
| ANXA10 | GGA2 | NAMPT | SHISA5 |
| APAF1 | GGT5 | NCOA7 | SLBP |
| ASPG | GPR160 | NFKB1 | SLC25A29 |
| ATXN3 | HCRTR2 | NLRC5 | SLC6A15 |
| B2M | HELZ2 | NMI | SMPD4 |
| BATF3 | HRH1 | NR2E1 | SNX3 |
| BCL2L15 | IFI6 | NT5C1B | SNX6 |
| BIRC2 | IFIH1 | NT5C3B | SOCS1 |
| BRWD1 | IFIT5 | NUB1 | STARD3NL |
| C1R | IFNAR2 | OASL | STAT1 |
| C1S | IFNG | OGFR | STOML1 |
| C26H6ORF222 | IGF2BP3 | OPTN | STX17 |
| C3AR1 | IL13RA1 | OR4S2 | SYNE1 |
| CADM1 | IL4I1 | OR5AS1 | TAP1 |
| CALCOCO2 | INF2 | ORC1 | TAP2 |
| CARS | IP6K2 | OTUD4 | TDRD6 |
| CCL19 | IRF1 | P2RX4 | TDRD7 |
| CCL5 | IRF2 | PARP12 | THEMIS2 |
| CCNA2 | IRF7 | PARP14 | TLDC2 |
| CEPT1 | KLHL10 | PARP9 | TLR3 |
| CMPK2 | L3MBTL3 | PFKP | TMPRSS2 |
| CMTR1 | LARP4B | PIGA | TNFAIP2 |
| CNP | LITAF | PLA2G6 | TOR2A |
| COL14A1 | LOC100857563 | PLCL2 | TRAF2 |
| COL6A1 | LOC100857706 | PLOD1 | TRAFD1 |
| COL6A2 | LOC101747234 | PLVAP | TRANK1 |
| COL6A6 | LOC101747378 | POSTN | TRIM13 |
| COL9A3 | LOC101749880 | PSME4 | TRIM25 |
| COMMD8 | LOC101750701 | PTPLAD2 | TSPAN32 |
| CSF2RB | LOC107049082 | PXK | TSTA3 |
| CX3CL1 | LOC107049337 | RAB40B | TTC3 |
| DDA1 | LOC107053690 | RASGEF1B | UBA2 |
| DDX60 | LOC107053736 | RBM43 | URI1 |
| DHX58 | LOC417192 | RBMS1 | USP18 |
| DNAJB9 | LOC418666 | RMDN3 | USP25 |
| DNAJC13 | LOC420721 | RNF114 | VCAM1 |
| DNAJC7 | LOC422513 | RNF144B | WDFY1 |
| DNMBP | LOC424214 | RNF19B | ZC3HAV1 |
| DRAM1 | LOC769676 | RNF213 | ZCCHC11 |
| DRAM2 | LY6E | RP9 | ZCCHC2 |
| DSEL | LYST | RPA1 | ZDHHC2 |
| EFCAB3 | MAP3K14 | RSAD2 | ZNFX1 |
| ELAVL2 | MARCKSL1 | RSPRY1 | ZP3L1 |
| ENDOV |  |  |  |

**Supplemental Table S4:** Regulation of select immune gene families in quail, chicken and duck lung in comparison to controls

|  | **Quail** |  | **Chicken** |  | **Duck** |  |
| --- | --- | --- | --- | --- | --- | --- |
|  | LPAI-1dpi | HPAI-3pi | LPAI-1dpi | HPAI-3dpi | LPAI-1dpi | HPAI-3pi |
| **IFITIM** |  |  |  |  |  |  |
| IFITM1 | ↑ 2 |  |  |  | ↑ 3 | ↑ 5 |
| IFITM2 |  |  | ↑ 2 |  | ↑ 4 | ↑ 6 |
| IFITM3 |  |  |  |  |  | ↑ 32 |
| **MHC I** |  |  |  |  |  |  |
| MHCI-1 | ↑ 2 | ↓ 5 |  |  |  | ↑ 2 |
| MHCI-2 | ↑ 2 | ↓ 5 |  |  |  | ↑ 7 |
| MHCI-3 |  |  |  |  |  | ↑ 3 |
| MHCI-4 |  |  |  |  | ↑ 2 | ↑ 2 |
| TAP1 | ↑ 2 | ↓ 3 |  |  |  |  |
| TAP2 | ↑ 2 | ↓ 4 |  |  |  |  |
| **MHC II** |  |  |  |  |  |  |
| MHCIIB-1 | ↑ 3 | ↓ 13 |  |  |  |  |
| MHCIIB-2 | ↑ 4 | ↓ 8 |  |  |  |  |
| MHCIIB-3 |  | ↓ 11 |  |  |  |  |
| MHCIIA | ↑ 2 | ↓ 18 |  |  |  |  |
| DMA | ↑ 4 | ↓ 10 |  |  |  |  |
| DMB1 | ↑ 3 | ↓ 15 |  |  |  |  |
| **TLR** |  |  |  |  |  |  |
| TLR2B |  | ↓ 9 |  |  |  |  |
| TLR3 | ↑ 3 |  |  |  |  |  |
| TLR4 | ↑ 4 | ↓ 10 |  |  |  | ↑ 5 |
| TLR5 |  |  |  |  | ↑ 3 | ↑ 2 |
| TLR7 | ↑ 3 | ↓ 12 |  |  |  | ↑ 2 |

**Supplemental File S1**

List of unannotated quail genes and their manual annotation

**Supplemental File S2**

Location of breakpoints between chicken and quail chromosomes

**Supplemental File S3**

BED file containing location of ERVs in quail genome

**Supplemental File S4**

BED file containing location of ERVs in turkey genome

**Supplemental File S5**

Sequences of cathelicidins and defensin genes identified in quail genome

**Supplemental File S6**

List of selection signatures detected from the HSR / LSR lines

**Supplemental File S7**

Statistics from photoperiod differential expression study

**Supplemental File S8**

Pathway analysis from photoperiod study

**Supplemental File S9**

List of differentially expressed genes, FDR < 0.05 and fold change > 1.6 in infection study

**Supplemental File S10**

Overrepresented GO terms in differentially expressed genes in infection study

**Supplemental File S11**

Significant upregulation and downregulation of IFITM, MHC and TLR genes
